# A computational model of shared fine-scale structure in the human connectome

**DOI:** 10.1101/108738

**Authors:** J. Swaroop Guntupalli, Ma Feilong, James V. Haxby

## Abstract

Variation in cortical connectivity profiles is typically modeled as having a coarse spatial scale parcellated into interconnected brain areas. We created a high-dimensional common model of the human connectome to search for fine-scale structure that is shared across brains. Projecting individual connectivity data into this new common model connectome accounts for substantially more variance in the human connectome than do previous models. This newly discovered shared structure is closely related to fine-scale distinctions in representations of information. These results reveal a shared fine-scale structure that is a major component of the human connectome that coexists with coarse-scale, areal structure. This shared fine-scale structure was not captured in previous models and was, therefore, inaccessible to analysis and study.

**Author Summary:** Resting state fMRI has become a ubiquitous tool for measuring connectivity in normal and diseased brains. Current dominant models of connectivity are based on coarse-scale connectivity among brain regions, ignoring fine-scale structure within those regions. We developed a high-dimensional common model of the human connectome that captures both coarse and fine-scale structure of connectivity shared across brains. We showed that this shared fine-scale structure is related to fine-scale distinctions in representation of information, and our model accounts for substantially more shared variance of connectivity compared to previous models. Our model opens new territory — shared fine-scale structure, a dominant but mostly unexplored component of the human connectome — for analysis and study.

## Introduction

Resting state functional magnetic resonance imaging (rsfMRI) reveals patterns of functional connectivity that are used to investigate the human connectome [1-3] and parcellate the brain into interconnected areas that form brain systems and can be modeled as networks [4-11]. The connectivity of a single area is considered to be relatively homogeneous and typically is modeled as a mean connectivity profile. Cortical topography, however, has both a coarse scale of cortical areas and a finer scale of multiplexed topographies within areas [12-16]. Fine-scale within-area topographies are reflected in patterns of activity that can be measured with fMRI and decoded using multivariate pattern analysis (MVPA)[12,13,17]. Fine-scale variation in connectivity, however, has been overlooked due to poor anatomical alignment of this variation across individual brains. We ask here whether local variation in functional connectivity also has a fine-scale structure, similar to fine-scale response tuning topographies, and whether such variation can be captured in a common model with basis functions that are shared across brains.

We developed a new algorithm, connectivity hyperalignment (CHA), to model local variation in connectivity profiles with shared basis functions for connectivity profiles across individuals and individual-specific local topographies of those connectivity basis functions (Fig 1). The resultant common model connectome consists of transformation matrices for each individual brain, which contain individual-specific topographic basis functions, and a common model connectome space, which contains shared connectivity profiles (Fig 2). Individual transformation matrices transform an individual brain’s connectome, in its native anatomical coordinate space, into the common model space [13,16]. The individual transformation matrices and common model connectivity matrix are derived iteratively from training data. Validity testing is done on connectivity profiles and other functional parameters from independent test data that are hyperaligned into the common model connectome space. The results show that CHA can derive these shared basis functions from functional connectivity derived from neural activity while watching an audiovisual movie and from neural activity in the resting state.

**Fig 1.**
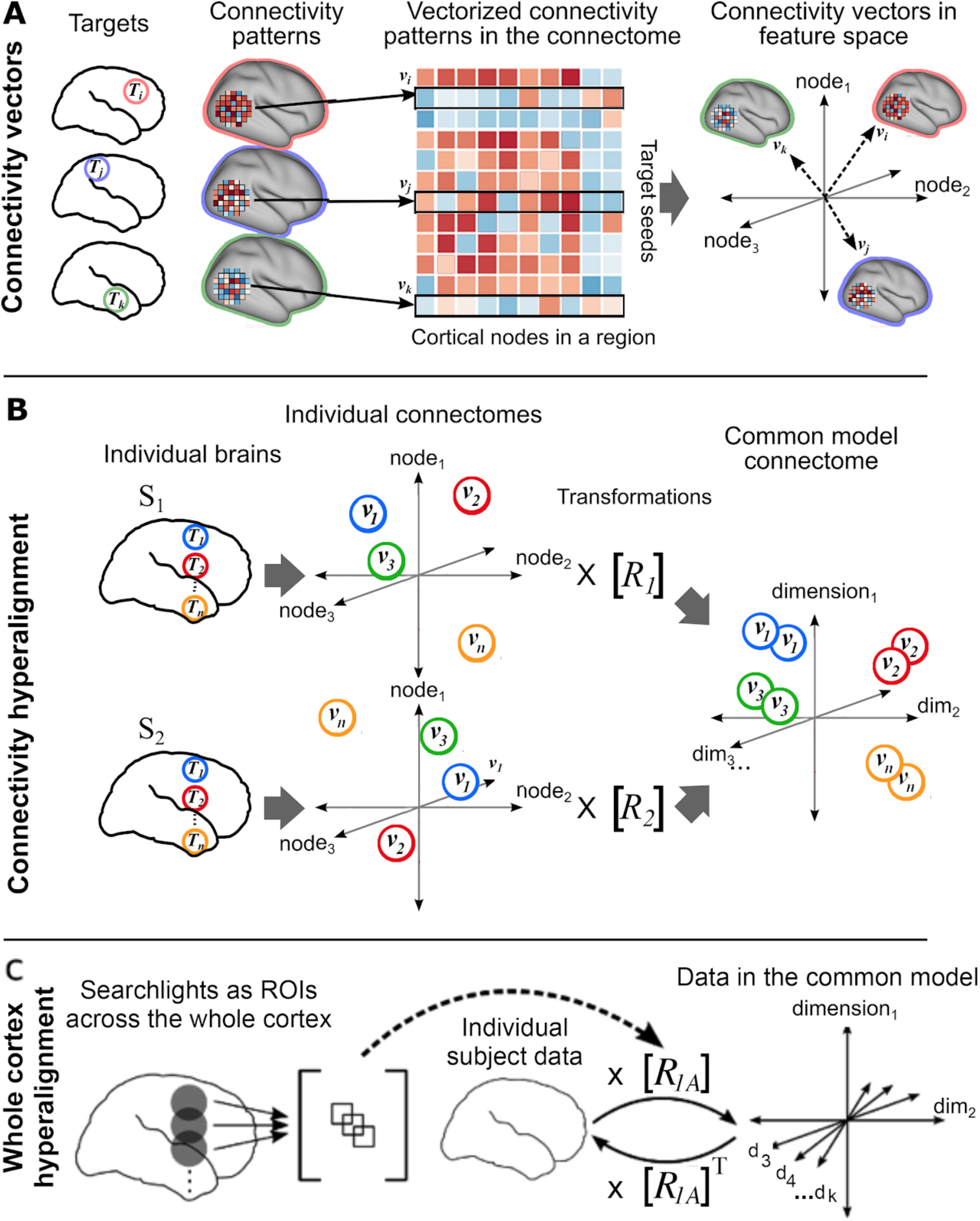
Schematic of connectivity hyperalignment (CHA). (A) Connectivity can be defined as any measure of similarity between a cortical locus (e.g., surface node/voxel) and a target region. Connectivities to a target region (***T***_i_, ***T***_j_, ***T***_k,_ …) of loci in a searchlight yield a connectivity pattern for that target in that searchlight. These patterns can be analyzed as connectivity pattern vectors (***v***_i_, ***v***_j_, ***v***_k,_, …) in a space in which each cortical locus in that region is a dimension. (B) Connectivity pattern vectors (***v***_1_, ***v***_2_, … ***v***_n_) in a region of interest or a searchlight to be hyperaligned are calculated for target regions (***T***_1_***, T***_2_…, ***T***_n,_) distributed uniformly across the whole cortex. At this stage connectivity hyperalignment derives transformation matrices for each brain (***R***_1_***, R***_2_, …) in each searchlight that align these vectors across subjects into a common high-dimensional connectivity space. (C) For each subject, searchlight transformation matrices are aggregated into a whole cortex transformation matrix, ***R***_1A_, as in [16], affording projection of connectivity data into a whole cortex common model connectome space. Conversely, the transpose of a whole cortex transformation matrix can project connectivity data from the whole cortex common connectome space back into that subject’s cortical anatomy.

**Figure 2.**
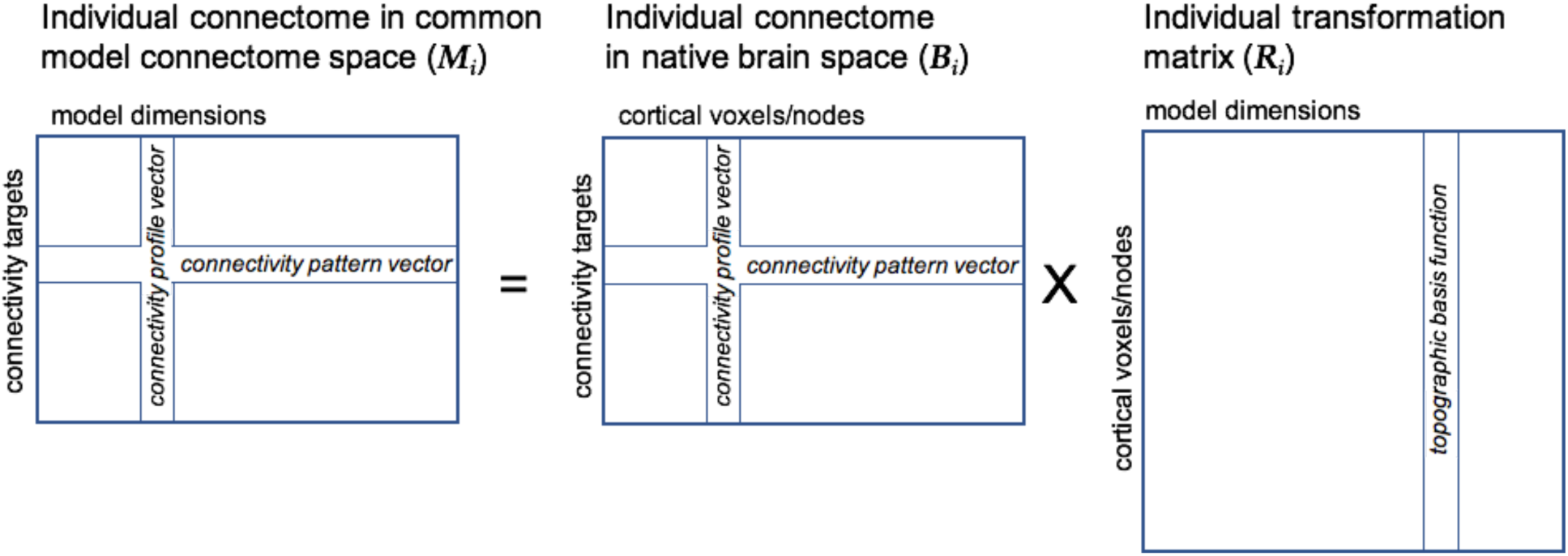
Schematic of data and transformation matrices for the common connectome. The connectivity data for an individual subject, ***i***, in that subject’s native brain space, ***B_i_***, is projected into the common model connectome space, ***M_i_**,* by multiplying it with the transformation matrix, ***R***_*i*_. Vectors in data matrix rows are connectivity pattern vectors — patterns of connectivity with a single connectivity target time-series across cortical nodes/voxels in the individual’s native brain space or across model dimensions in the common model connectome. Vectors in data matrix columns are connectivity profile vectors — connectivities of a single node/voxel or model dimension across connectivity targets. The transformation matrix contains weights for the linear transformation of connectivity vectors in an individual’s brain data space into the common model connectome space. Vectors in transformation matrix columns for model dimensions are patterns of weights for a local field of voxels/nodes and serve as topographic basis functions. Individual variation in the fine-scale topographic pattern of connectivity to a target is modeled as a weighted mixture of multiplexed or overlaid topographies for model dimensions.

The resultant common model connectome accounts for substantially more shared variance in functional connectivity derived from both movie fMRI data and resting state fMRI data than was accounted for by previous models. This shared variance resides in fine-scale local variations in connectivity. We show further that this local variability in functional connectivity profiles is meaningful in that it is closely related to local patterns of response that encode fine distinctions among representations. Our results indicate that shared fine-scale local variation, which was not evident in previous models, is a major component of the human connectome that coexists with shared coarse-scale areal structure. Our common model connectome makes this fine-scale local variation accessible for group-level study of its network properties.

## Results

We derived a common model of the human connectome by applying CHA to fMRI data collected while 11 subjects viewed a full-length movie [13,16] and to rsfMRI data for 20 subjects in the Human Connectome Project (HCP) database [18-20]. The common model connectome is high-dimensional with connectivity profiles for model dimensions that serve as basis functions for modeling the connectivity profiles of cortical loci in individual brains. We validated the common model in terms of 1) increased intersubject correlations (ISCs) of connectivity profiles, and 2) increased spatial specificity of shared connectivity profiles. To test whether this fine-scale structure is meaningful for the representation of information, we tested the effect of CHA on 3) ISCs of representational geometry for the movie, 4) between-subject multivariate pattern classification (bsMVPC) of responses to the movie and 5) ISCs of task activation and contrast maps from the HCP database. The first two validation experiments are designed to test whether connectivity hyperalignment improves alignment of functional connectivity across brains in a way that preserves the fine-grain spatial granularity of variation in connectivity profiles. These validations were tested on functional connectivity derived from both the movie and rsfMRI data. The third, fourth, and fifth validation experiments are designed to test whether the transformation of individual brain spaces into the common model space better aligns topographies associated with representation of information and cognitive processes. The third and fourth validations were tested on the movie data. The fifth validation test used rsfMRI and task fMRI data from the HCP database.

### Intersubject correlation of connectivity profiles

CHA afforded large increases in ISCs of connectivity profile vectors in both the movie fMRI data and the rsfMRI data (Figures 3 and 4).

**Fig 3.**
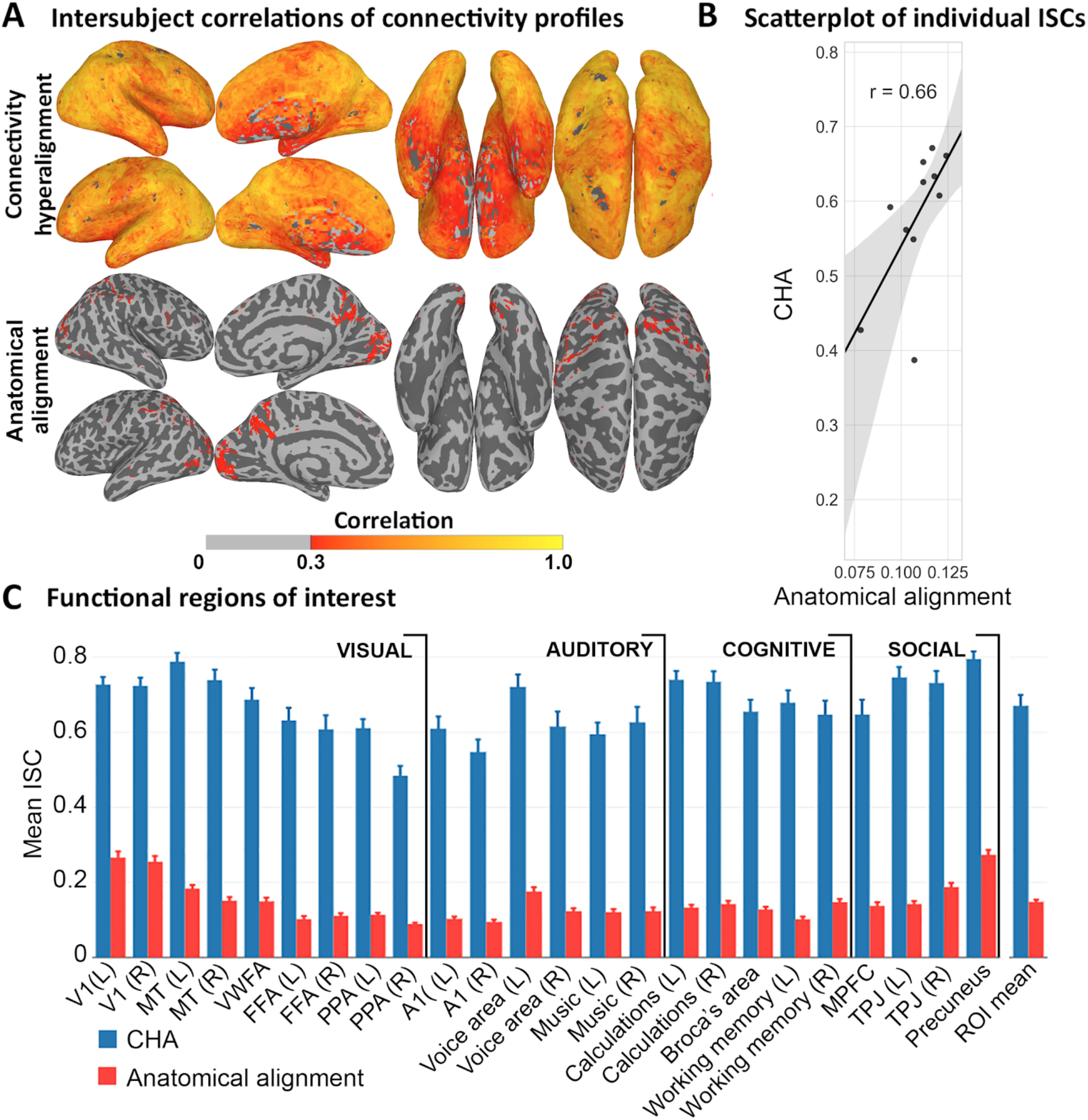
ISC of connectivity profiles calculated from movie data. (A) Average ISCs of connectivity profiles in each surface node after CHA and anatomical alignment. (B) Scatter plot of individual whole cortex mean ISCs of connectivity profiles before and after CHA with linear fit. Each subject’s similarity of connectome with the group is improved by CHA while preserving similarity or deviance from others. Shaded region is the 95% CI. (C) Mean ISCs of connectivity profiles in functional ROIs covering visual, auditory, cognitive, and social systems comparing the common model connectome space and anatomical alignment. Bootstrapped testing showed significantly higher ISCs after CHA than after anatomical alignment in all ROIs.

**Fig 4.**
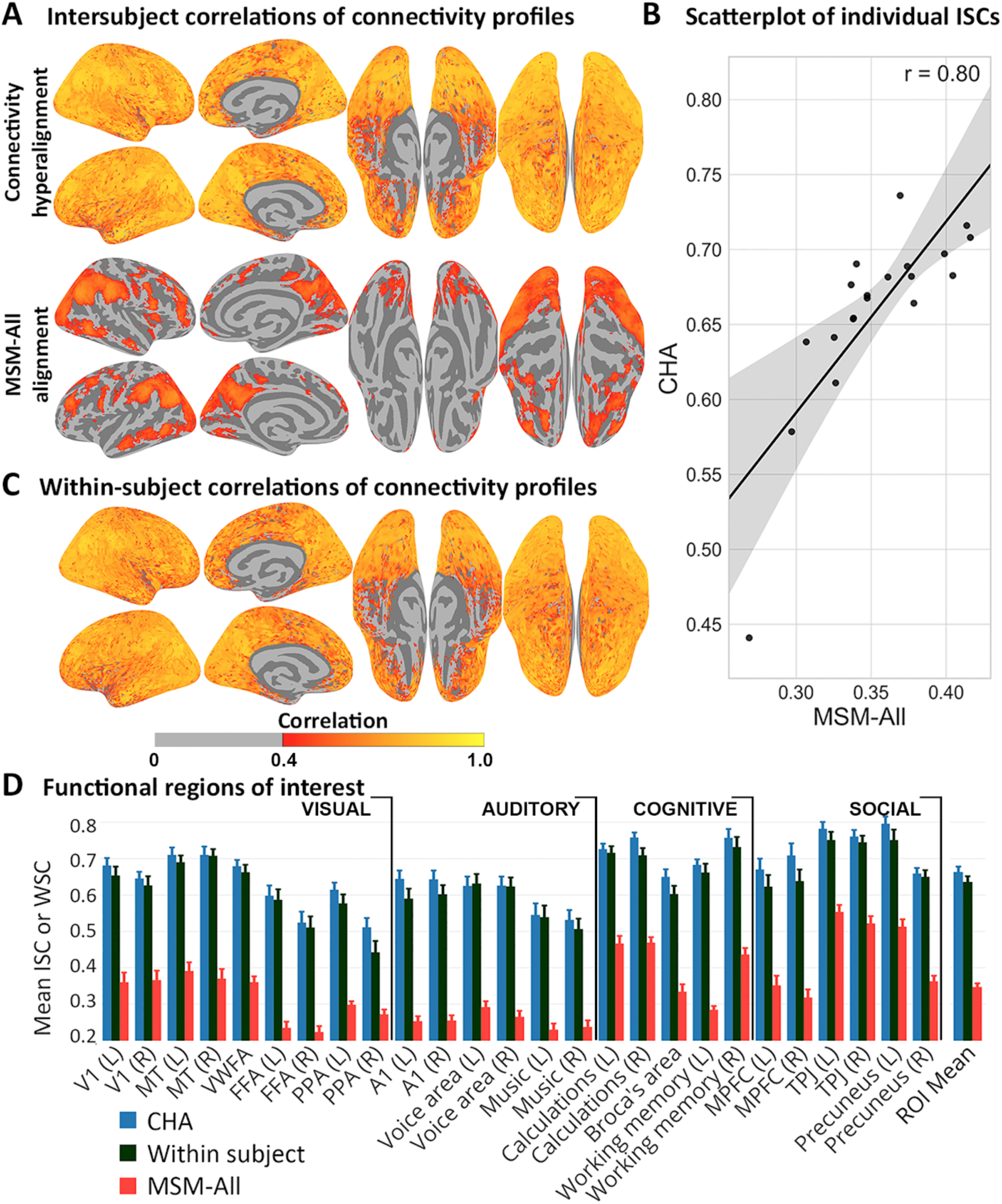
ISC of connectivity profiles calculated from HCP rsfMRI data. (A) Average ISC of connectivity profiles in each surface node in the common model connectome space and after surface alignment (MSM-All). (B) Scatter plot of individual whole cortex mean ISCs of connectivity profiles before and after CHA with linear fit. Each subject’s similarity of connectome with the group is improved by CHA while preserving similarity or deviance from others. Shaded region is the 95% CI. (C) Average within-subject between-session correlations in the common space. (D) Mean ISCs and WSCs of connectivity profiles in functional ROIs covering visual, auditory, cognitive, and social systems comparing the common model connectome space, within-subject between-session correlation in common space, and surface alignment.

Increases in ISCs of functional connectivity derived from movie data were distributed across all of cortex (Fig 3). ISC at a cortical node is the correlation of the one subject’s connectivity profile with the mean of other subjects’ profiles, indexing how well other subjects’ connectivity profiles can predict an individual’s connectivity profiles. Fig 3A shows a cortical map of mean ISCs of connectivity profiles in the common model connectome space as compared to ISCs in anatomically-aligned data. Fig 3B is a scatterplot of mean ISCs for individuals after anatomical alignment and CHA, which shows that CHA increased ISC for each individual and preserved individual similarity or deviance from the group. We quantify the increases in 24 functional ROIs, identified using a meta-analytic database, NeuroSynth [21](Fig 3C; table S1). Mean ISC of connectivity profiles across these ROIs was markedly higher in the common model connectome than in the anatomically-aligned data (0.67 versus 0.15; difference = 0.52, 95% confidence interval, CI = [0.46, 0.56]).

Increases in ISCs of resting state connectivity profiles were similarly distributed across all of cortex and replicated the findings based on ISCs of movie viewing connectivity profiles (Fig 4). Fig 4A shows a cortical map of mean ISCs of resting state connectivity profiles in the common model connectome space and in data aligned with the HCP’s MSM-All method (multimodal surface matching [22]). Fig 4B is a scatterplot of mean ISCs for individuals after MSM-All alignment and CHA, which shows that CHA of resting state data increased ISC for each individual and preserved individual similarity or deviance from the group. Fig 4C shows a cortical map of within-subject correlations between connectivity profiles from different resting state sessions. We quantify the increases in 26 functional ROIs, identified using a meta-analytic database, NeuroSynth [21](Fig 4D; table S1). Mean ISC of connectivity profiles across these ROIs was markedly higher in the common model connectome than in the MSM-All-aligned data (0.66 versus 0.35; difference = 0.31 [0.30, 0.33]). ISCs of resting state connectivity profiles in the common model connectome space are slightly higher than within-subject correlations of resting state connectivity profiles (mean correlation = 0.64; CI for difference = [0.00, 0.05]) (Fig 4D). This latter result indicates that an individual’s connectome based on resting state functional connectivity is better predicted by the common model connectome, based on other subjects’ data, than by estimates based on a typical sample of that subject’s own rsfMRI data, due to the benefit of estimating connectivity profiles based on a large number of brains and the precision of CHA.

The substantial increase in ISCs with hyperalignment is due in part to discovery of shared variance that was obscured by misalignment but also to suppression of unshared variance and amplification of shared variance mediated by filtering the data in the transformation step with smaller weights for nodes with unshared or noisy variance and larger weights for nodes with shared variance. To gauge the size of the effect of filtering independent of better information alignment, we calculated ISCs in data that are filtered by our algorithm but aligned based on anatomy or MSM-All (see methods). ROI mean ISCs of connectivity profiles in movie data filtered with CHA but aligned based on anatomy was 0.22 (CHA versus filter-control difference = 0.45 [0.39 0.49]) and for HCP resting state data filtered with CHA but aligned based on MSM-All was 0.41 (CHA versus filter-control difference = 0.25 [0.23, 0.27]). These ISCs are larger than ISCs of unfiltered, anatomically and MSM-All-aligned data but, nonetheless, still markedly lower than ISCs of connectivity profiles in the common model connectome space, which is both filtered and re-aligned by CHA.

### Spatial granularity of connectivity profile variation

We investigated the spatial specificity of the common model connectome by computing the intersubject spatial point spread functions (PSF) of ISCs of connectivity profiles [16]. The PSF of connectivity profiles was computed as the correlation of the connectivity profile in a cortical surface node for a given subject with the average connectivity profiles of other subjects in the same node and nodes at cortical distances ranging from 3 to 12 mm. We similarly calculated within-subject PSFs based on within-subject correlations (WSC) of connectivity profiles between two resting state sessions. Fig 5A shows the slopes of connectivity profile PSFs for movie data in 24 functionally-defined ROIs, and Fig 5B shows the mean PSF across these ROIs as a function of cortical distance) in the common model connectome space and in anatomically-aligned data. CHA increased the average slope of PSF across these ROIs, relative to anatomical alignment, from 0.013 to 0.105 (difference=0.092 [0.080, 0.099]). Fig 5C shows the slopes of connectivity profile PSFs for resting state connectivity profiles in the 26 functionally-defined ROIs, and Fig 5D shows the mean PSF across these ROIs (ISC or WSC as a function of cortical distance) in the common model connectome space, in MSM-All-aligned data, and within-subject. CHA increased the average slope of PSF across these ROIs, relative to MSM-All alignment, from 0.012 to 0.065 (difference=0.053 [0.047, 0.055]). The intersubject PSF slopes in the common model connectome space and the PSF within-subject (slope = 0.067) were not significantly different (difference = 0.002 [−0.002, 0.007]). This fine spatial granularity was ubiquitous in cortex, with steep PSFs in sensory-perceptual areas in occipital and temporal cortices as well as in higher-order cognitive areas in lateral and medial parietal and prefrontal cortices.

**Fig 5.**
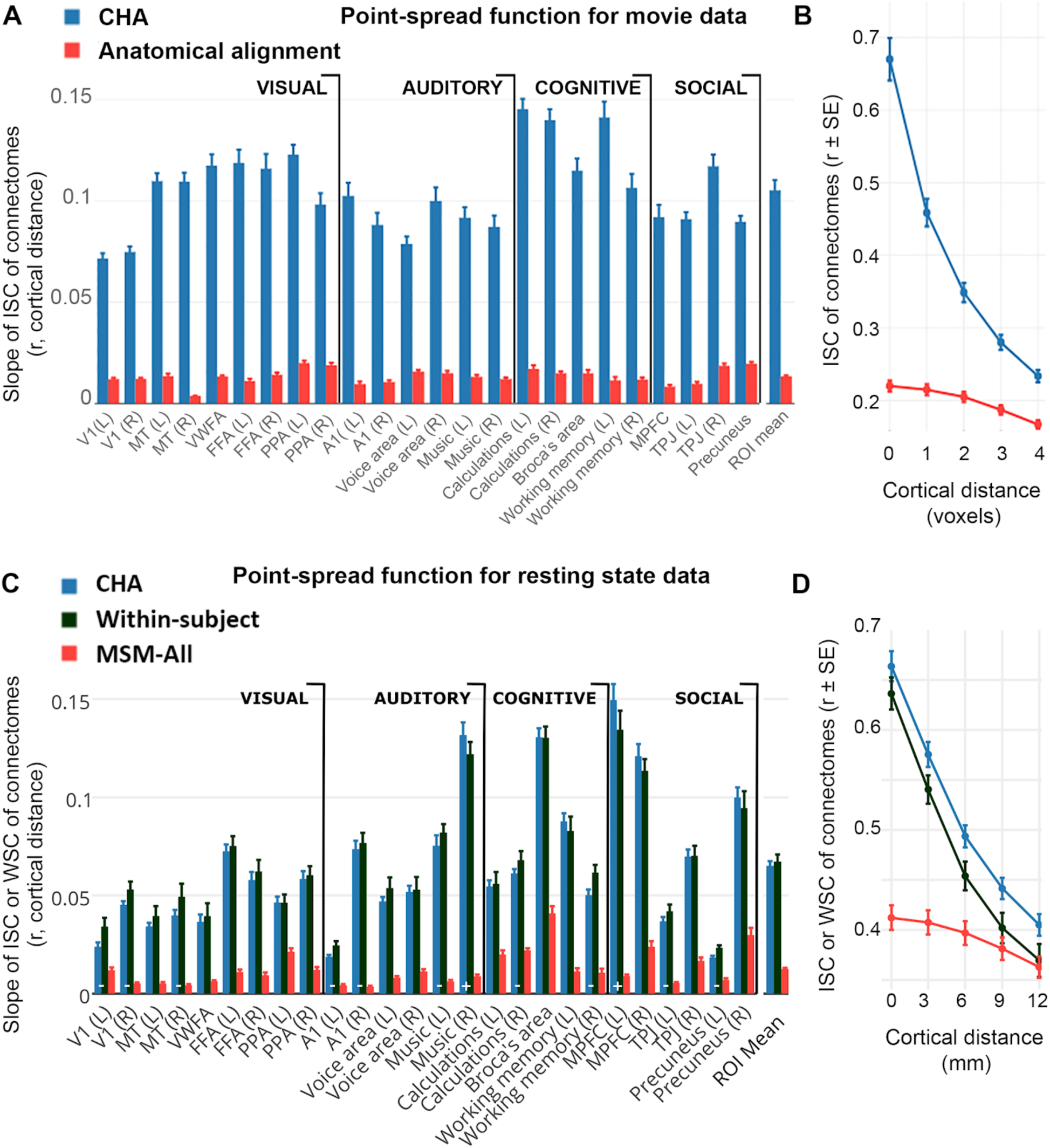
Spatial granularity of shared connectivity profiles. The intersubject point spread function (PSF) of connectivity profile correlations are computed as the correlation between the connectivity profile for a cortical locus in one subject and the profiles of the same locus and its spatial neighbors in other subjects at increasing distances from that locus. For the HCP rsfMRI data, within-subject PSFs are computed as the correlation between the connectivity profile for a cortical locus from one rsfMRI session and the profiles of the same locus and its spatial neighbors from a different rsfMRI session. Slope is estimated in each functional ROI as the linear fit of intersubject or within subject correlations as a function of distance. (A) Slope of PSFs for movie viewing connectivity profiles in 24 functional ROIs. (B) Average movie viewing connectivity PSF across all ROIs is plotted as ISC as a function of cortical distance. (C) Slope of PSFs for resting state connectivity profiles in 26 functional ROIs. (D) Average resting state connectivity PSF across all ROIs is plotted as ISC or WSC as a function of cortical distance.

The mean PSFs across ROIs (Fig 5B and 5D), clearly show that CHA captures fine-scale variations in connectivity profiles for neighboring cortical nodes across subjects that are not captured by anatomical alignment or MSM-All alignment. The ISCs of connectivity profiles for neighboring nodes in the common model connectome are substantially lower than ISCs for the same node (movie data: 0.21 [0.18, 0.23]; resting state: 0.09 [0.08,0.09]). Similar fine spatial granularity is seen in the within-subject between-session PSFs for resting state connectivity profiles (0.10 [0.09, 0.10]). By contrast, ISCs for connectivity profiles in the anatomically-aligned and MSM-All aligned data barely differ for nodes spaced 0 versus 1 voxel/3 mm (differences =0.005, [0.005, 0.005] and 0.004, [0.004,0.015], respectively) and 2 voxels/6 mm (0.01 [0.01, 0.02] and 0.02 [0.01, 0.02], respectively) apart. Decrements for larger distances (ISCs of nodes spaced 3 voxels/9 mm: 0.03 [0.03, 0.03] and 0.03 [0.03,0.03], respectively; and 4 voxels/12 mm: (0.05 [0.05, 0.06] and 0.05 [0.04,0.05], respectively) were similarly small.

### Generalization to fine-scale patterns in response tuning

Next we asked if this shared variance in fine-scale local variation in connectivity profiles carries meaning by testing whether it reflects fine-scale variations in response tuning topographies that carry fine-grained distinctions in representation. We tested whether projecting movie response data into the CHA-derived common connectome space afforded better alignment of representational geometry for movie time-points and better bsMVPC of movie time segments.

Results show that shared fine-scale structure in the common model connectome is closely related to fine distinctions in representations. Fig 6A shows a cortical map of mean ISCs of local representational geometry after anatomical alignment and in the common model connectome. Representational geometry is the matrix of all pairwise similarities between patterns of response to different time-points in the movie, resulting in a matrix of more than 800,000 pairwise similarities (see methods). Fig 6B shows a cortical map of mean bsMVPC accuracies for 15 s movie time-segments in searchlights after anatomical alignment and CHA. CHA greatly increased both ISCs of representational geometry and bsMVPC accuracies. Quantification of these effects in functional ROIs is illustrated in Fig S3. CHA significantly increased ISCs of representational geometry in all ROIs (ROI mean ISCs = 0.308 and 0.210 after CHA and anatomical alignment, respectively, difference = 0.097 [0.080, 0.110]). CHA also dramatically increased bsMVPC accuracies in all ROIs (ROI mean bsMVPC accuracies = 10.37% and 1.04% after CHA and anatomical alignment, respectively, difference = 9.33% [7.71%, 10.54%]).

**Fig 6.**
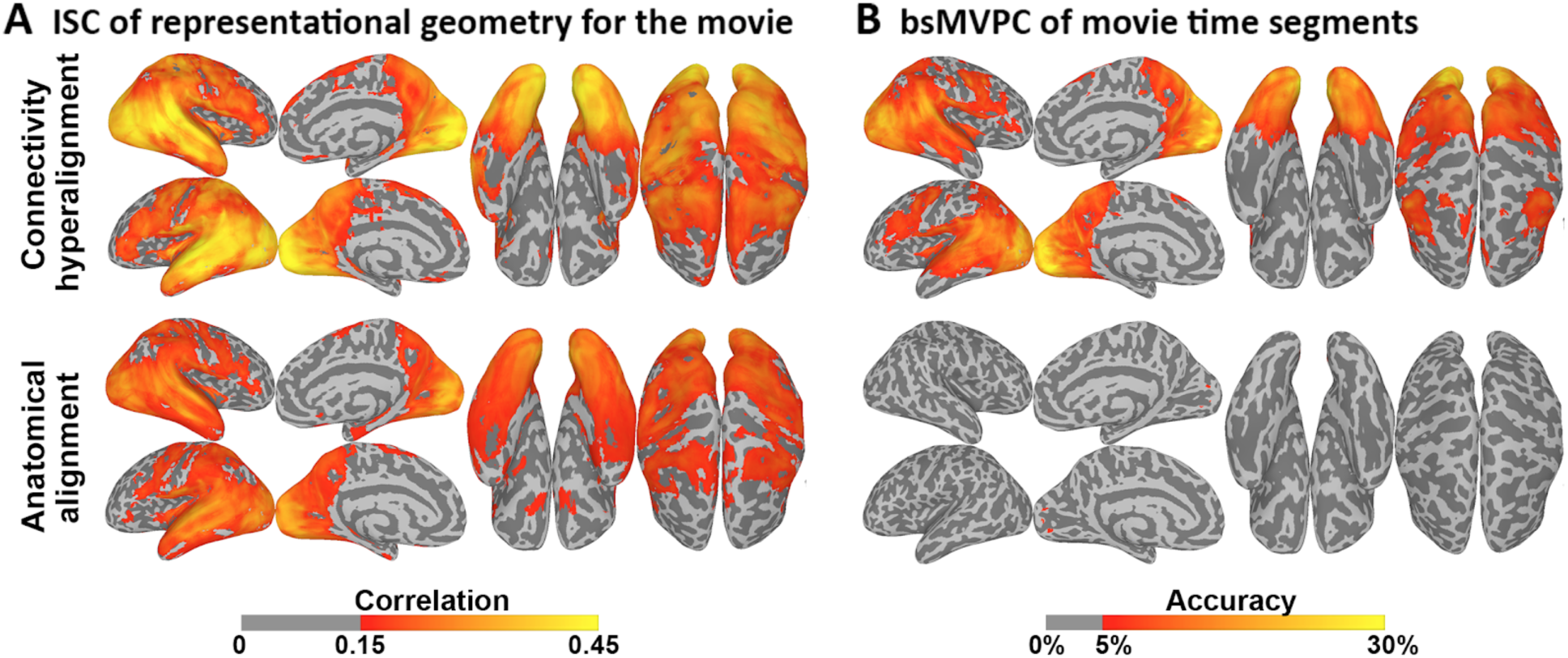
Effect of CHA on ISC or representational geometries and bsMVPC of movie data. (A) ISC of representational geometry in each voxel mapped onto the cortical surface. (B) Accuracies for bsMVPC of 15 s movie segments. Classification was performed within each movie half separately, and the accuracies are then averaged across the two halves. Parameters for hyperalignment are derived from the half that was not used for classification.

### Generalization to task maps from the HCP database

We tested the generalization of the common model connectome derived from resting state fMRI by applying connectivity hyperalignment parameters derived from one session of resting state data to task maps provided by the HCP database comprised of 32 task activation maps and 14 task contrast maps (Supplemental Table S2). These task maps reflect simple operations and, thus, do not have the same fine-grained structure that is associated with activation by dynamic, naturalistic stimuli such as a movie. We calculated the ISC of these task maps between each subject and the average of others before and after hyperalignment. Hyperalignment improved correlations on average across all tasks and in all but two (Face-Shapes and Body-Average, labeled ns) task contrast maps (Fig 7). The average correlation across task maps increased from 0.58 to 0.65 (mean difference = 0.07 [0.06, 0.08]).

**Figure 7.**
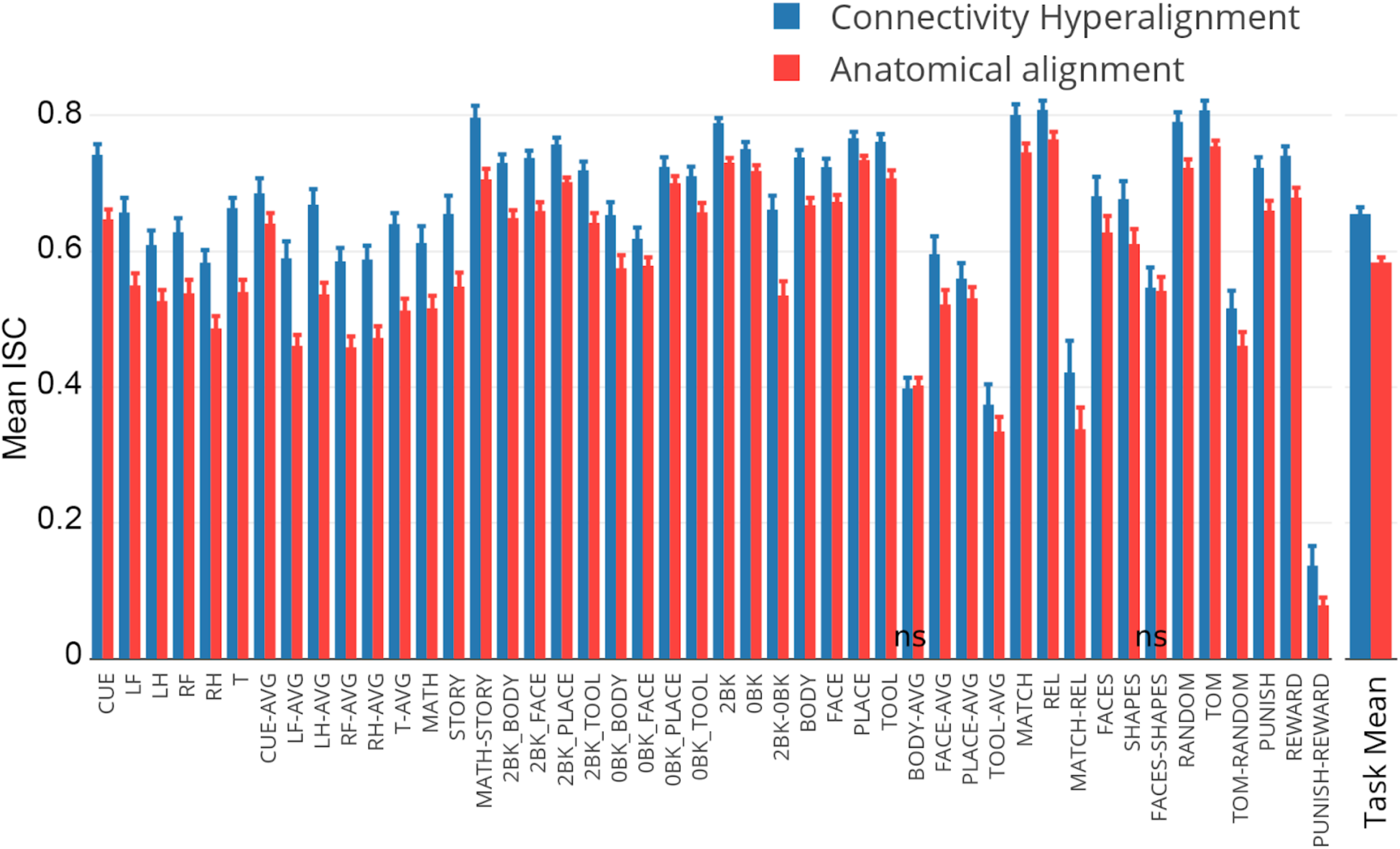
ISCs of HCP task activation and contrast maps after CHA and surface alignment (MSM-All).

### Comparison of CHA and Response Hyperalignment (RHA)

Since CHA aligned fine-scale patterns of response tuning functions across subjects better than anatomy-based alignment, we asked how well it compares to our previously published response-based hyperalignment (RHA) [16]. Because RHA requires responses that are synchronized across subjects in time, it cannot be applied to resting state data. We compare CHA and RHA of movie viewing data on 1) ISC of connectivity profiles, 2) ISC of representational geometry, and 3) bsMVPC of 15 s movie segments.

Results showed that both CHA and RHA increased ISCs and bsMVPC classification accuracies significantly over anatomy-based alignment, but each algorithm achieves better alignment for the information that it uses to derive a common model, namely connectivity profiles and patterns of response, respectively. ISCs of connectivity profiles are significantly higher in a common model based on CHA than in a common model based on RHA (ROI mean ISCs = 0.67 and 0.575, respectively; CHA-RHA difference = 0.095 [0.081, 0.112])(Supplemental Figure S2). By contrast, RHA marginally but significantly outperforms CHA on some validations based on response tuning functions, namely ISCs of representational geometry (ROI means = 0.322 and 0.308, respectively; RHA-CHA difference = 0.014 [0.007, 0.019])(Supplemental Figure S3), and bsMVPC of movie segments (ROI mean accuracies = 13.65% and 10.37%, respectively; RHA-CHA difference = 3.28% [2.76%, 3.78%])(Supplemental Figure S4).

## Discussion

These results show that fine-scale local variation in connectivity profile is a major component of the human connectome that can be modeled with shared connectivity basis functions. Each connectivity basis function has a connectivity profile that is shared across subjects and a different local connectivity topography in each individual brain. These basis functions are derived from multiple subject data in local cortical fields. An individual’s connectivity pattern in a cortical field is modeled as multiplexed or overlaid connectivity topographic basis functions, and the connectivity profile of each cortical node or voxel is modeled as a weighted mixture of local connectivity profile basis functions. Thus, the connectivity profile for each voxel or node is modeled as a high-dimensional vector of connectivity profile bases, capturing how it varies locally from its neighbors, rather than modeling the connectivity of a brain area as a single connectivity profile that is shared by all voxels or nodes. We show that these shared basis functions can be discovered with connectivity hyperalignment of data collected during viewing and listening to a rich naturalistic movie and during the resting state. These basis functions constitute a common model connectome. Shared fine-scale variation is a ubiquitous characteristic of all of human cortex and is a major component of the human connectome that coexists with shared coarse-scale areal variation.

We show that patterns of connectivity exhibit fine-scale variation that is captured in the CHA-derived common model connectome. We define fine-scale structure as voxel-by-voxel or node-by-node variation in response and connectivity profiles, as compared to the coarse structure of parcels that consist of sets of voxels or surface nodes and are treated as a functional unit with a homogeneous functional profile. In Figure 8 we illustrate the fine scale structure that is captured in the common model connectome for connectivity patterns in a left lateral-occipital/inferior-temporal cortex cortical field. Quantitatively, we show that shared fine-scale structure is captured in the common model connectome with a direct measure of the spatial granularity of local variation in connectivity profiles — the intersubject point-spread function. The intersubject spatial point-spread function for variation in connectivity profiles is dramatically, six to eight-fold, steeper after data are transformed into the common model connectome than for data that are anatomically aligned. Next we show that capture of this fine-scale structure in functional connectivity generalizes to capture of fine-scale structure in neural representation. Transformation of movie data into the common model space, using matrices derived from functional connectivity in independent movie data, afford bsMVPC of time segments that are tenfold higher than for anatomically-aligned data. bsMVPC of movie time segments relies on fine-scale structure that is not well-aligned based on anatomy, nor on functional alignment using a “rubber-sheet” warping of cortical topographies, nor on hyperalignment based on responses to a limited variety of still images of visual categories [13,16, 23-25]. ISCs of local representational geometries also are dramatically higher after CHA than after anatomical alignment. These local representational geometries reflect fine-scale structure that reveals how information spaces in different cortical fields vary, offering a window on how these spaces are transformed along processing pathways and reshaped by task demands [26-28]. Finally, we also show that transformations derived from rsfMRI improve alignment of topographies in task activation and task contrast maps in the HCP database.

**Figure 8.**
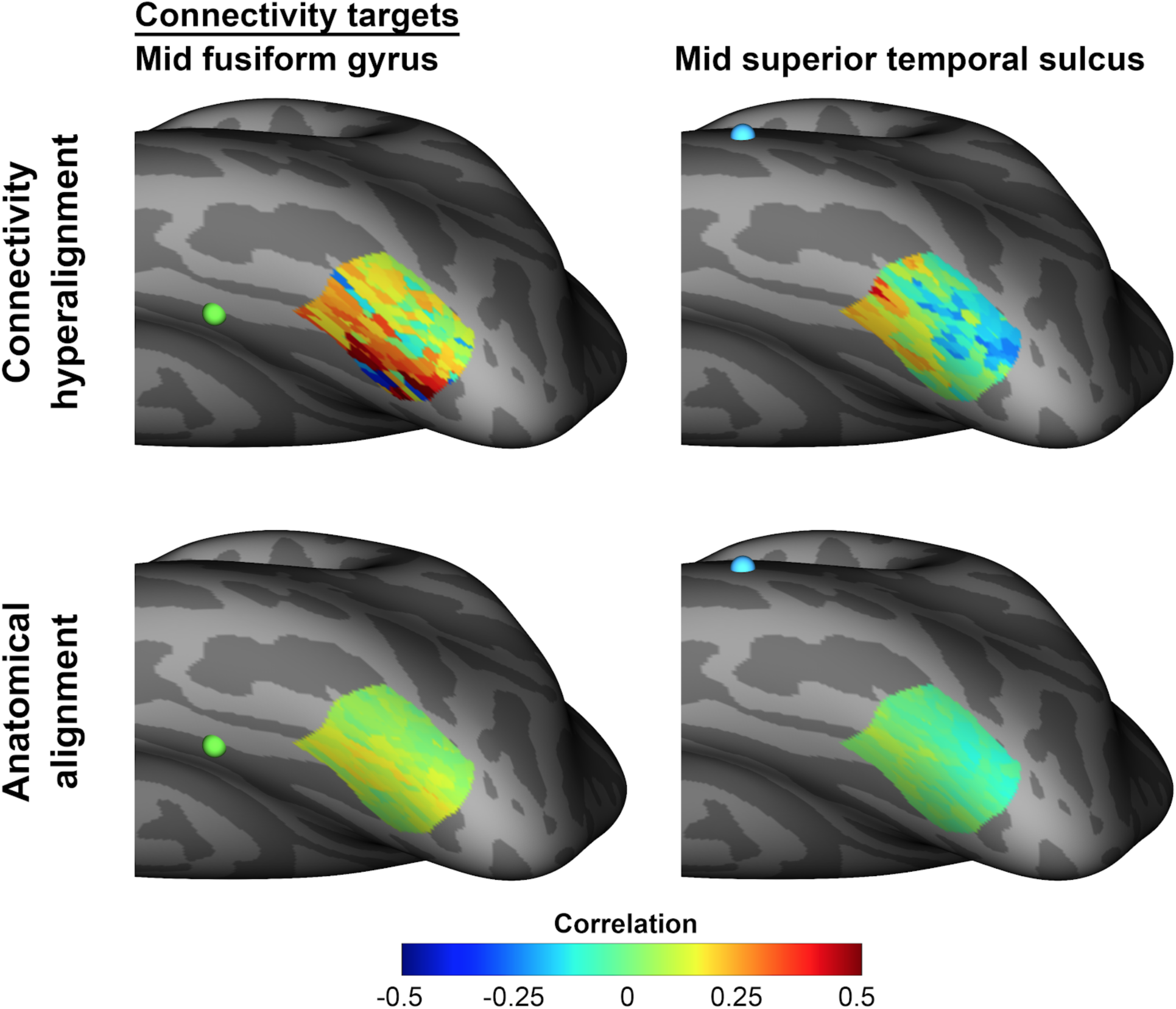
Mean group connectivity patterns in a left lateral-occipital/inferior temporal cortical field. Connectivity patterns were measured from movie data for functional connectivity with connectivity targets in mid lateral fusiform gyrus and mid superior temporal sulcus. Mean group connectivity patterns are shown for data in the common model connectome, derived with CHA based on responses to the other half of the movie, and for anatomically aligned data. Mean ISCs for patterns after CHA are higher than after anatomical alignment for both the fusiform target (0.835 versus 0.175) and the STS target (0.826 versus 0.306). The occipitotemporal, mid fusiform, and mid STS loci are taken from the face-responsive fields identified by Visconti di Oleggio Castello, Halchenko, et al. [28]. The locations of the fusiform and STS targets are indicated with green and blue dots, respectively. The inflated cortical surface is tipped to provide a clear view of the cortical field. Connectivities are correlations of time-series responses to the movie.

The existence and importance of fine-scale connectivity is well-recognized [29-31] but previously was not modeled in a common computational framework and, consequently, was largely overlooked. Attempts to model within-area topographies of connectivity either were limited mostly to within-subject analyses or coarser within-area topographies that could be captured with anatomy-based alignment of group data [31]. Consequently, when not simply overlooked, within-area variations in connectivity profiles were usually analyzed as gradients that have a single cycle in a cortical area, such as retinotopy or somatotopy [29-31].

Other models of shared structure in the human connectome have focused on the identification of shared functional networks that can be identified with cluster analysis (e.g. [5,32,33]) or independent components analysis (ICA; e.g. [19]). These methods do not attempt to align the fine-scale structure within areas in these networks. In some approaches, each voxel is assigned to one cluster or system and is, thereby, associated with the time-series tuning function that characterizes that cluster [5,8,32,33]. Approaches that use ICA, or related componential analyses such as PCA or SVD, have the potential to capture node-by-node variation in connectivity profiles, but implementations of these approaches have not adapted them to analyze this fine-scale topographic structure. For example, dual regression could allow using group ICA as a common space for modeling each voxel in an individual as a weighted sum of independent components [34,35]. In practice, however, each voxel is characterized in ICA analyses by the network to which it belongs, not as a mixture of multiplexed functional topographies. Node-by-node local variation in connectivity topographies is blurred in group analyses because individual variation on independent components is projected into anatomically-aligned brains rather than into a single reference voxel space to reveal shared fine-scale structure, as we do here. A novel approach by Langs et al. [32,33] allows nodes to be assigned to different clusters in a common functional connectivity embedding space independently of anatomical location. The implementations of this method, however, do not attempt to discover shared fine-scale structure, and the low dimensionality of the embedding space and small number of clusters are probably insufficient to capture this level of detail.

Cortical functional architecture has multiplexed topographies at multiple spatial scales. In primary visual cortex, retinotopy is multiplexed with ocular dominance columns, edge orientation, spatial frequency, motion direction, and motion velocity, among other low-level visual attributes [36,37]. Primary visual cortex sends coherent projections to other visual areas where these topographies are recapitulated and transformed, affording the emergence of more complex features, such as curvature, texture, shape, color constancy, and biological motion; and, subsequently, even higher-order attributes such as object categories, view-invariant face identity, and species-invariant attributes of animals such as action categories and dangerousness [12,15,26-28,38-40]. Similar transformations of multiplexed topographies characterize other sensory modalities and, undoubtedly, supramodal cognitive operations. Modeling inter-areal communication as a single value of connectivity strength sheds no light on how information is transformed along cortical processing pathways to allow high-order information to be disentangled from confounding attributes [41].

Multiplexed cortical topographies at multiple spatial scales can be modeled with individual-specific topographic basis functions that have shared tuning profiles [13,16] and shared connectivity profiles (as shown here). No previous model captured multiple spatial scales of connectivity topographies with connectivity profiles that are shared across brains. By capturing coarse- and fine-scale connectivity topographies with shared basis functions, the common model connectome casts a bright light on the dominant role of fine-scale connectivity patterns in the human connectome and opens new territory for investigation of the network properties of cortical connectivity at finer levels of detail. With this new perspective, inter-areal connectivity can be modeled as more than a simple replication of global activity, as is the assumption underlying existing approaches to modeling the connectome, but, instead, as information processing operations in which functional topographies are transformed by projections between areas.

## Methods

### Movie data: Raiders of the Lost Ark

We scanned 11 healthy young right-handed participants (4 females; Mean age: 24.6+/-3.7 years) during movie viewing. Participants had no history of neurological or psychiatric illness. All had normal or corrected-to-normal vision. Informed consent was collected in accordance with the procedures set by the local Committee for the Protection of Human Subjects. Participants were paid for their participation. These data also were used in a prior publication on whole cortex RHA [16].

#### Stimuli and design

Stimuli consisted of the full-length feature movie — “Raiders of the Lost Ark” — divided into eight parts of approximately 14 to 15 min duration. Video was projected onto a rear projection screen with an LCD projector which the subject viewed through a mirror on the head coil. The video image subtended a visual angle of approximately 22.7° horizontally and 17° vertically. Audio was presented through MR Confon’s MRI-compatible headphones. Participants were instructed to pay attention to the movie and enjoy. See [16] for details.

#### fMRI protocol

Participants were scanned in a Philips Intera Achieva 3T scanner with an 8 channel head coil at the Dartmouth Brain Imaging Center. T1-weighted anatomical scans were acquired at the end of each session (MPRAGE, TR=9.85 s, TE=4.53 s, flip angle=8°, 256 × 256 matrix, FOV=240 mm, 160 1 mm thick sagittal slices). The voxel resolution was 0.9375 mm × 0.9375 mm × 1.0 mm. Functional scans of the whole brain were acquired with an echo planar imaging sequence (TR=2.5 s, TE=35 ms, flip angle=90°, 80 × 80 matrix, FOV=240 mm × 240 mm) every 2.5 s with whole brain coverage (41 3 mm thick interleaved axial slices, giving isotropic 3 mm × 3 mm × 3 mm voxels). We acquired a total of 2718 functional scans with 1350 TRs in four runs during the first session and 1368 TRs in four runs during the second session.

#### fMRI data preprocessing

fMRI movie data were preprocessed using AFNI software [42](http://afni.nimh.nih.gov). Functional data were first corrected for the order of slice acquisition and head motion by aligning to the last volume of the last functional run. Any spikes in the data were removed using 3dDespike in AFNI. Data were then filtered using 3dBandpass in AFNI to remove any temporal signal variation slower than 0.00667 Hz, faster than 0.1 Hz or that correlated with the whole brain average signal or the head movement parameters. Each subject’s anatomical volume was first aligned to the motion corrected average EPI volume and then to the MNI 152 brain template in AFNI. Functional EPI BOLD data were then aligned to the MNI 152 brain template using nearest neighbor resampling by applying the transformation derived from the alignment of the anatomical volume to the template. Data acquired during the overlapping movie segments were discarded resulting in a total of 2662 TRs with 1326 TRs in the first session and 1336 TRs in the second session.

#### Definition of masks and searchlights for movie data

We derived a gray matter mask by segmenting the MNI_avg152T1 brain provided in AFNI and removing any voxel that was outside the cortical surface by more than twice the thickness of the gray matter at each surface node. It included 54,034 3 mm isotropic voxels across both hemispheres. We used this mask for all subsequent analyses of all subjects.

Hyperalignment of movie data started with hyperalignment of data in 20,484 overlapping searchlights of 20 mm radius centered on cortical nodes with 2.9 mm average spacing between the nodes. Cortical nodes were defined in a standard cortical surface from FreeSurfer (fsaverage)(https://surfer.nmr.mgh.harvard.edu) and resampled into a regular grid using AFNI’s MapIcosahedron [42,43] with 10,242 nodes in each hemisphere. We defined the surface searchlights [44] in PyMVPA [45](http://www.pymvpa.org) as cortical disks. The thickness of disks was extended beyond the gray matter, as defined in FreeSurfer, 1.5 times inside the white-matter gray-matter boundary and 1.0 times outside the gray-matter pial surface boundary to accommodate any misalignment of gray matter as computed from the anatomical scan and the gray matter voxels in the EPI scan. To reduce the contribution from noisy or non-gray matter voxels that were included due to this dilation, we used a between-subject correlation measure on training data [13] to select 70% of the voxels in each searchlight [16]. The mean number of selected voxels in movie data searchlights was 235.

Searchlights for defining connectivity targets were defined using a coarse surface grid corresponding to the ico8 surface in SUMA [43] with 1284 nodes (10.7 mm spacing between nodes). We used surface disk searchlights [44] centered on these nodes as the movie data connectivity target searchlights. These searchlights had a radius of 13 mm, as did those used for the HCP data, producing complete coverage of the cortex with overlapping searchlights. Cortical disks centered on these voxels were dilated using the same procedure as for hyperalignment of cortical surface searchlights. Movie connectivity target searchlights had a mean of 99 voxels.

### Resting state data: Human Connectome Project

In the HCP database [20], we found unrelated subjects of age <= 35 with at least four resting state scans, yielding a list of 64 subjects. We chose the first 20 of these subjects in the sorted order of subject IDs for our analysis.

For each subject, we used their cortical surfaces and fMRI data aligned to the group using MSM-All [22] with 32K nodes in each hemisphere as provided by the HCP. We used data from one resting state session [19](“rfMRI_REST1_LR”) to derive CHA parameters and validated it on a different resting state session (“rfMRI_REST2_LR”), and task fMRI sessions [18](EMOTION, GAMBLING, LANGUAGE, MOTOR, RELATIONAL, SOCIAL, and WM). Resting state data were acquired for 1200 TRs with a TR of 0.720s in each session (total time=14 min 33 s). The data used to derive the CHA parameters and common model and the resting state data used for validation tests used the same phase-encoding direction (LR). We used a single session of rsfMRI for alignment to mimic a typical resting state data acquisition which usually varies from 10-20 mins of scanning. See [19] for more details about the acquisition and preprocessing pipelines.

#### Definition of masks and searchlights for HCP data

We masked the data to include only the left and right cortices (Cortex_Left and Cortex_Right), removing all the non-zero nodes that correspond to the medial subcortical regions, resulting in 59,412 nodes across both hemispheres. These nodes also defined the centers of 59,412 surface searchlights [44] with 20 mm radii that were used for hyperalignment. All nodes in these searchlights were included. The mean number of surface nodes in the HCP searchlights was 337.

We defined connectivity target searchlights using a coarser surface grid corresponding to the ico8 surface in SUMA [43] with 1284 nodes (10.7 mm spacing between nodes). We found the closest matching nodes on the 32K surface to the nodes on the ico8 surface, and used those as centers for connectivity target searchlights. These searchlights had a radius of 13 mm, producing complete coverage of the cortex with overlapping searchlights. HCP connectivity target searchlights had a mean of 142 loci. See further details below for how time-series were extracted from these searchlights.

For validation of task fMRI, we used all of the maps provided by the HCP after removing redundancies (such as FACE-AVG and AVG-FACE), which resulted in 46 maps (Supplemental Table S2).

### Connectivity Hyperalignment

We use CHA to derive a common model of the human connectome and the transformation matrices that project individual brains’ connectomes into the common model connectome space. The common model connectome is a high-dimensional information space. In the current implementation, the model space based on movie fMRI data has 54,034 dimensions, corresponding to the number of voxels in the gray matter mask, and the model space based on HCP resting state fMRI data has 59,412 dimensions, corresponding to the number of cortical nodes in those data. The derivation of this space starts with hyperalignment in local cortical fields, searchlights, which yields orthogonal transformation matrices for each subject in each field. These searchlights are aligned across subjects based on anatomy (movie data) or MSM-All (HCP resting state data); consequently, each locus within a searchlight is similarly aligned across subjects before CHA. Local transformation matrices for each searchlight map anatomically or MSM-All aligned cortical loci in a cortical field to CHA-aligned dimensions in the common model connectome. These local transformation matrices are then aggregated into a whole brain transformation matrix, which is not globally orthogonal. The whole brain transformation matrices are derived based on local hyperalignment in searchlights to constrain resampling of information to cortical neighborhoods defined by those searchlights.

#### The basic equation for hyperalignment (both CHA and RHA)

*B*_*ij*_ are the original matrices of data for cortical fields, *j*, in individual brains, *i,* which have *m*_*ij*_ columns of cortical loci and *n* rows of data vectors. Hyperalignment derives a transformation matrix for each cortical field in each individual, *R*_*ij*_, and a matrix for each cortical field, *M*_*j*_, that is the mean of transformed individual brain matrices, *B_ij_R_ij_*, minimizing the Frobenius norm of differences between transformed individual brain matrices and the model space matrix. For each cortical field *j*:

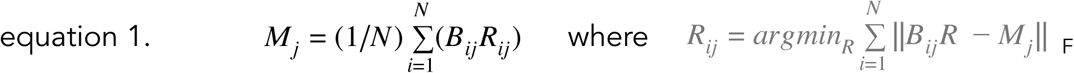

For whole cortex hyperalignment we define the cortical fields, *j*, as searchlights. Thus, we estimate a transformation matrix, *R*_*ij*_, for each of *N*_*sl*_ searchlights in each subject *i*. We then aggregate these searchlight transformation matrices into a whole cortex transformation matrix, *R*_*iA*_ (details below):

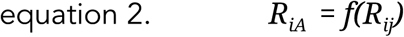

The whole cortex common model data matrix, *M*, is created by transforming individual whole cortex data matrices, *B_iA_,* into common model space coordinates and calculating the mean:

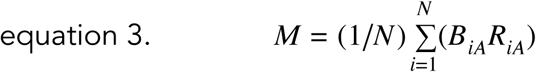

Conversely, other subjects’ data in the common model space can be mapped into any subject’s individual anatomical space using the transpose of that subject’s whole cortex transformation matrix, 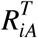, producing a data matrix, *M*_*i*_, in which the columns are that subject’s cortical loci, making it possible to analyze and visualize transformed group data in any subject’s anatomical space:

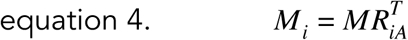

In our implementations of hyperalignment, we have used a variant of Generalized Procrustes Analysis [46,47](described in detail below) to derive orthogonal transformation matrices for the improper rotations of a brain data matrix from a cortical field (region of interest or searchlight) to the mean of others’ matrices for the same region to minimize interindividual differences between the transformed individual and mean data matrices. Aggregation of searchlight transformation matrices, *R*_*ij*_, produces a whole cortex transformation matrix, *R*_*iA*_. Because *R*_*iA*_ is derived from searchlight transformation matrices, *R*_*ij*_, it imposes a locality constraint that limits remapping of brain data to nearby cortical loci (see details below), making the whole cortex transformation matrix nonorthogonal by design. We also have tested other hyperalignment algorithms that use alternatives for calculating the transformation matrices, such as regularized canonical correlation and probabilistic estimation [48,49]. These alternatives are effective but have not yet been extended to aggregate local transformation matrices for cortical fields into a whole cortex transformation matrix.

The dimensionality of the brain and model data matrices is *n* × *m*, in which *m* equals the number of cortical nodes or dimensions in brain and model data matrices — *B*_*ij*_, *B*_*iA*_, *M*_*j*_, and *M* — and *n* equals the number of data vectors across these dimensions. The number of data vectors, *n*, is set and determined by the number of connectivity targets for defining connectivity pattern vectors (see details below). For RHA, *n* is set by the number of response pattern vectors in an experimental dataset. The number of cortical loci in a cortical field or searchlight, *m*_*ij*_, can vary across subjects. If the number of cortical loci or dimensions differs between subjects or between an individual subject and the model space, the new subject’s data are transformed into a space with the same dimensionality as the first subject’s or the model’s space. The number of cortical loci in the whole cortex model is set at *m* = 59,412 for HCP data and *m* = 54,034 for movie data. We also have shown that the dimensionality of a model for region of interest or searchlight (*m*_*j*_) can be reduced substantially relative to the dimensionality of individual brain spaces in imaging datasets (*m*_*ij*_), *m_Mj_ << m_ij_*[13,16,48]. In the current version of CHA, as in whole cortex RHA, however, we do not reduce the dimensionality of the model space because these reduced dimensionality local models are difficult to aggregate into a whole cortex model.

Note that the common model data matrix has two distinct components. The columns define a common model space, whereas the rows are defined by the experimental data — either patterns of connectivity to targets elsewhere in the brain for CHA, or patterns of response for RHA. The space can be illustrated as an anatomical space insofar as it can be rotated into any individual’s cortical loci (equation 4), but there is no “canonical” anatomical space, rather the individuality of each individual brain is preserved. We illustrate results in the anatomical space of one subject, the “reference subject”, but we also could illustrate the results in other subjects’ anatomical spaces. The special nature of the common space derives from the alignment of functional indices — connectivities and responses — to minimize interindividual differences and, thereby, discover shared basis functions for the individually variable functional architecture. These basis functions are the response and connectivity profiles for model dimensions that model the response and connectivity profiles of cortical loci in individual brains as linear weighted sums. In other words, equation 4 models single columns in *B*_*i*_ as weighted sums of columns in *M*_*i*_.

Transformation matrices consist only of weights for the projection of individual brain spaces for cortical fields, *B*_*ij*_, or the whole cortex, *B*_*iA*_, into model spaces (*M_j_, M*) and contain no connectivity or response data. Thus, a transformation matrix can be applied to any matrix of data vectors in an individual brain space. Similarly, the transposes of transformation matrices, 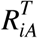, can be applied to any data vector in the model space to project that vector into the cortical topographies of individual brains. For all applications of the common model, including the validation tests presented here, the transformation matrices are applied to independent data that played no role in derivation of the model space and the individual transformation matrix parameters. This is necessary to avoid overfitting [50]. Transformation matrices derived from connectivity data also can be applied to response data and vice versa. In other words, RHA and CHA are complementary methods for deriving a common model of information spaces in cortex, and RHA-derived and CHA-derived transformation matrices are alternative projections for mapping individual brain data into the same common model space. Note that each column of the transformation matrix *R*_*iA*_ contains weights for cortical loci in subject *i*’s brain. These columns of weights are basis functions for modeling functional topographies in individual brains as linear weighted sums of topographies associated with model dimension functional profiles.

#### Derivation of transformation matrices for regions of interest and searchlights

The derivation of individual transformation matrices that map individual brain spaces into the common model space is a three-level iterative process. We present the iterative algorithm for deriving transformation matrices and the common model space in greater detail here to help readers understand better its structure.

In the first step of the first level, the data matrix for a cortical field in one subject, ***B**_2j_*, is transformed to be in optimal alignment with the same cortical field in another subject’s brain, ***B**_1j_*, referred to here as the reference subject:

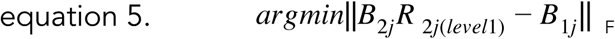

We use the Procrustes transformation to find the orthogonal matrix that affords the optimal improper rotation to achieve this minimization [46]. Note that this “rotation” is a rotation of data in the high-dimensional feature space, not a rotation in a two or three dimensional anatomical space. Elsewhere we have shown that other algorithms can be used to achieve this minimization [48,49].

In the following steps of the first level, the brain data matrices for the third and subsequent subjects are transformed to be in optimal alignment with the matrix defined by the mean of the previous subject’s matrix and the previous mean:

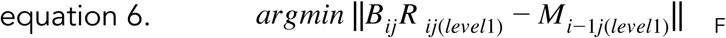

where

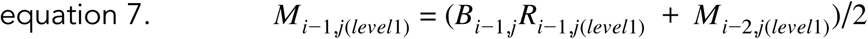

***M***_*i-1,j(level1)*_ is the target data used to hyperalign the current subject’s data, ***B***_*ij*_, and ***M***_*i-2,j(level1)*_ is the target data used to hyperalign the previous subject’s data, ***B**_i-1,j_*. Target data is updated with previous subjects’ aligned data in this first level. In the subsequent two levels each subject’s data matrix is hyperaligned to the simple, unweighted mean of all other subjects’ matrices.

At the end of the first level, level one transformation matrices have been derived for all cortical fields in all subjects, ***R**_i,j(level1)_,* which are used to project each subjects’ brain data into the provisional common spaces that evolved over level one iterations ***M**_i,j(level1)_*. Each subject is then re-hyperaligned to the mean data matrix for all other subjects’ transformed data from level one to derive new individual transformation matrices, ***R**_ij(level2)_*. Note that the new transformation matrices are derived using each subject’s original brain data, ***B***_*ij*_. Note also that the mean matrices in provisional common spaces, *M* _¬*i*,*j*(*level*1)_, exclude data from the subject being hyperaligned:

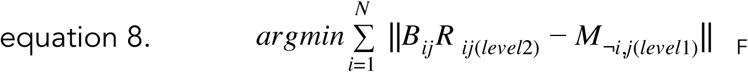

where

*M* _¬*i*,*j*(*level*1)_ is the equally-weighted mean of level one transformed data for all subjects but subject *i*:

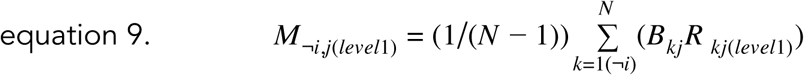

After the level two transformation matrices, ***R**_ij(level2)_*, are calculated for each subject, the level one transformation matrices are discarded, and the group mean of transformed individual brain data matrices is recalculated, using these new transformation matrices, producing the model matrix, ***M***:

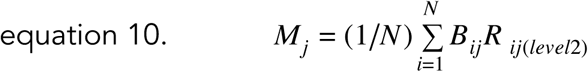

In level 3, the last level, the final searchlight transformation matrices, ***R***_*ij*_, are recalculated for each subject (see equation 1 above).

#### Derivation of whole cortex transformation matrices

Orthogonal transformation matrices for hyperaligning a cortical field can map information from a cortical locus into model dimensions anywhere else in that cortical field. To constrain the remapping of information to nearby locations in the reference subject’s cortical anatomy, we developed a searchlight-based approach [16]. We hyperalign the data in ***N***_*sl*_ overlapping searchlights, where ***N***_*sl*_ is the number of searchlights (59,412 for HCP data, 20,484 for movie data). The number of model dimensions in each searchlight transformation matrix is determined in the movie data by cortical location and the number of selected features in the reference subject (mean = 235) and in HCP data by the cortical location of the searchlight (mean = 337). The transformation for each searchlight, ***R***_*ij*_, has dimensionality that corresponds to the number of nodes in an individual’s searchlight (***m***_*ij*_ rows) and the number model dimensions in that searchlight, derived from the reference subject’s anatomy (***m***_*1j*_ rows). Thus, each transformation matrix has on the order of 28K and 57K free parameters for movie data and HCP data, respectively. Because the searchlights are overlapping, there are multiple estimates of weights for mapping each cortical locus to each model dimension. As described in Guntupalli et al. [16] these weights are aggregated across searchlights by adding all weights for each cortical-locus-to-model-dimension mapping. In essence, this is equivalent to creating a whole cortex transformation matrix, ***R***_*iA*_, of dimensionality *m × m*, by padding each searchlight transformation matrix, ***R***_*ij*_, with zeroes in all rows and columns for cortical loci and model dimensions that are not in the individual or model searchlight cortical field to give them the same dimensionality to produce ***R**_ij(padded)_*, and then summing these padded transformation matrices. Thus, for each subject, *i*:

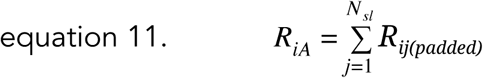

where ***N***_*sl*_ is the number of searchlights and ***R***_*ij(padded)*_ is the padded transformation matrix for subject *i* in searchlight *j* with dimensionality *m × m*. As noted above, *m* is the number of cortical loci — 59,412 surface nodes for HCP data and 54,034 voxels in the gray matter mask for movie data — and the number of whole cortex model dimensions. Because the searchlight approach constrains cortical-locus-to-model-dimension mapping to nearby cortical locations, the whole cortex transformation matrix, ***R***_*iA*_, is sparse with zero weights for all mappings of cortical loci to model dimensions that are separated by more than 2x the searchlight diameter (~4 cm in this implementation). The whole cortex transformation matrices, ***R***_*iA*_, are large (*m × m*) but sparse. 98.7% of the entries are zeros, and roughly 20 million entries have nonzero values in each of these matrices. The additive aggregation of mapping parameters weights nearby cortical location pairs, which co-occur in more searchlights than distant pairs, more strongly than distant pairs, adding a further locality constraint. Note that the searchlight transformation matrices, ***R***_*ij*_, are orthogonal but the whole cortex transformation matrices, ***R***_*iA*_, are not by design, to introduce the locality constraint. The whole cortex transformation matrices, ***R***_*iA*_, are used in all validation tests to hyperalign independent new data matrices after normalizing the data in each cortical node or voxel. In other words, all validation tests are performed on independent data that played no role in deriving the transformation matrix parameters or the common model connectome, providing cross-validated generalization testing. CHA of movie data was based on one half of the movie data (~55 min, ~1300 TRs) and the other, independent half of the movie data was used for validation tests with two-fold cross-validation. CHA of HCP data was based on one session of resting state data (~15 min, 1200 TRs) and a second session of independent resting state data was used for validation tests, as well as independent data from task fMRI [18].

Note that transformations map the cortical loci of a subject’s data matrices (columns) into the reference subject’s cortical loci. Thus, we use the reference subject’s cortex for illustration, but note that the anatomical coordinates for model dimensions are an abstraction, as even the reference subject’s data are mapped into model space coordinates with a transformation matrix that is not the identity matrix. Data matrices in the model space also can be mapped into any subject’s cortical anatomy by using the transpose of that subject’s transformation matrix (equation 4). Thus, the hyperaligned data in the common model space can be illustrated in any subject’s anatomical space. The anatomical space that we use for illustration, that of the reference subject, should not be considered a canonical space but, rather, simply as one of many possible physical instantiations.

#### Connectivity targets

We define functional connectivities as the correlations of the response profiles — series of responses across time — of cortical loci or dimensions with the response profiles of targets (***t***_*j*_) distributed across the cortex. We use two sets of connectivity targets, one reduced set to derive the transformation matrices and common model connectivity data matrix, and a more complete set to test the validity of the model. We define a reduced set of connectivity targets using surface-searchlight target ROIs to make derivation of the model more computationally tractable, as compared to using all cortical loci as individual connectivity targets. For the reduced set, we use 3852 targets (top 3 components for 1284 searchlights; note that the searchlights for connectivity targets are different from the searchlights that are hyperaligned as described above in the Resting State Data and Movie Data sections; see details for defining searchlight PC connectivity targets in the next section). For validation testing we analyze the full connectome, defining connectivity targets as all cortical loci in the brain (***N***_*cl*_ = 54,034 gray matter voxels in the movie data and 59,412 cortical nodes in the HCP resting state data).

#### Searchlight ROI connectivity targets

Each surface-searchlight connectivity target has a radius of 13 mm and is centered on a node of a coarse surface with a total of 1284 nodes covering both hemispheres. Thus, neighboring connectivity targets searchlights are overlapping. Unlike others (e.g., [5]) we do not assume that a searchlight connectivity target has a single response profile. We find, rather, a variety of response profiles for individual cortical loci in a target searchlight that can be captured as principal components. We used the top three principal components to represent the response profiles in a target searchlight.

To insure that the top components in target searchlights capture the same connectivity patterns across subjects, we performed a singular value decomposition (SVD) on the group mean connectivity matrix for each target searchlight after a simplified hyperalignment of individual matrices. Note that using a naive PCA/SVD to derive top components in each subject’s searchlight independently will not guarantee their functional similarity. Target searchlights had a mean of 142 loci (HCP data) or 99 voxels (movie data). At this stage it was not yet possible to break the response profiles for searchlight targets into multiple components with shared connectivity profiles. Consequently, connectivity targets for the procedure to derive these components were simply the mean time-series responses for target searchlights. For each target searchlight with ***N***_*s*_ features (surface nodes or voxels), we computed a 1284 × ***N***_*s*_ correlation matrix (the correlations between each cortical locus in the target searchlight and the mean time series for all target searchlights) for each subject. We hyperaligned the features (cortical loci) in each target searchlight across subjects based on these matrices and calculated the mean correlation matrix after hyperalignment in each target searchlight. We then performed a singular value decomposition (SVD) of each searchlight’s group mean matrix to obtain the top three components that explained the most shared variance. Each of these components is a weighted sum of cortical loci in a target searchlight for each subject, and these weights afford calculation of a time-series response whose connectivity profile with other targets is shared across subjects. Each individual subject’s time-series responses for the top three components were then used as target response profiles for CHA. This step gave us 1284 × 3 = 3852 target response profiles in each subject’s cortex.

### Validation tests and statistical analyses

#### Functional ROIs

In addition to analyzing the results of validation tests in each feature or searchlight across the whole cortex, we also examined the results of validation tests in functional ROIs associated with different sensory, perceptual, and cognitive functions to assess the general validity of the common model [16]. We searched for terms and cortical areas implicated in visual, auditory, cognitive, and social functions in NeuroSynth [22] and took the coordinates for the peak location associated with each of 24 terms (Supplementary Table 1). For validation testing using the movie dataset, we used volume searchlights centered around those peak loci with a radius of 3 voxels as our functional ROIs. For validation testing using the HCP dataset, we found the closest surface node corresponding to each peak locus and used a surface searchlight with a 10 mm radius around that surface node as the functional ROI. Functional ROIs that were medial and encompassing both hemispheres in the volume space were split into left and right ROIs in the surface space resulting in 26 ROIs for tests on the HCP data. For analyses of ISCs and PSFs of connectivity profiles in functional ROIs, we calculated the mean ISC or PSF across all cortical loci within the ROI searchlights (Figs. 3C, 4D, 5A, and 5C).

#### Statistics

We used bootstrapping to test for significance of the contrasts between alignment methods by sampling subjects 10,000 times to compute 95% CIs using BootES [51]. We did this for each ROI and for the mean of all ROIs separately. We used the same bootstrapping procedure for all validation tests unless specified otherwise.

#### Control for effect of filtering

In addition to the anatomically-aligned movie data and MSM-All aligned HCP resting state fMRI data, we calculated a third dataset that controls for the effect of filtering the data through CHA transformations but aligns those filtered data across subjects based on anatomical or MSM-All alignment. To produce the filter control data, we created multiple common model connectomes using each subject as the reference. Each subject’s connectome was transformed into the common connectome whose reference subject was the next subject in our order of subjects. The last subject’s connectome was transformed into the common model connectome whose reference brain was that of the first subject. Thus, each subject’s connectome is filtered by hyperalignment, but since the common model connectome for each subject has a different reference, the correspondence across filtered connectomes is based only on anatomical alignment and preserves the anatomical variability in the movie data and HCP datasets.

#### Intersubject correlation (ISC) of connectivity profile vectors

For validity testing we applied a more detailed definition of the connectome to measure fine-grained structure. The connectivity profile vector for a feature (or a cortical node or voxel) was defined as the correlation of its time-series with of all other cortical nodes or voxels. ISCs of connectivity profiles were computed between each subject’s connectivity profiles and the average connectivity profiles of all other subjects in each cortical locus.

For the movie data ISCs of connectivity profiles were computed within each movie half separately and before and after CHA based on the other half of the movie. Correlation values were Fisher transformed before averaging across both halves of the movie in each voxel. These were then averaged across all subjects and inverse Fisher transformed before mapping onto the cortical surface for visualization. ISCs of resting state connectivity profiles were computed for session REST2. Session REST1 was used for deriving the common model connectome and transformation matrices. ISCs were calculated for data mapped into the common model connectome, for movie data aligned anatomically, for HCP resting state data aligned with MSM-All, and for filter control movie and HCP data.

We also computed within-subject between-session (REST1 and REST2) correlation of resting state connectivity profiles. Within-subject between-sessions correlations were calculated on data that are transformed by CHA as used for our main analyses.

#### Spatial point spread function

To investigate the spatial granularity of representation, we computed a spatial point spread function (PSF) of ISCs or WSCs of connectivity profiles. We computed the correlation of connectivity profiles in each cortical locus (surface node or voxel) with the average connectivity profiles of cortical loci at varying cortical distances in other subjects’ data. To account for the effect of filtering, we did this analyses with filter control data that were filtered with CHA but aligned based on anatomy and MSM-All and after CHA with each subject aligned to the same reference subject [16]. We computed similar PSFs for connectivity profiles within-subject between-sessions (REST1 and REST2). This was also performed after CHA to account for any filtering effects but to a single common space as used for our main analyses.

#### ISC of representational geometry

ISCs of similarity structures were computed within each movie half separately using a searchlight of 3 voxel radius. The mean number of voxels in these searchlights was 102. In each searchlight, similarity structure was computed as a matrix of correlation coefficients between patterns of response for every pair of time-points from that movie half for each subject. The flattened upper triangle of this matrix excluding the diagonal was extracted as the profile of representational geometry at each searchlight for each subject. ISC of representational geometry in each searchlight was computed as the correlation between each subject’s representational geometry and the average of all other subjects’ representational geometries for that searchlight. Correlation values were Fisher transformed before averaging across both movie halves in each voxel. These were then averaged across all subject-average pairs and inverse Fisher transformed before mapping onto the cortical surface for visualization. The same steps were performed to compute inter-subject correlation of representational similarity before and after hyperalignment.

#### Between-subject multivariate pattern classification (bsMVPC)

bsMVPC of 15 s movie time segments (6 TRs) was computed within each movie half separately using searchlights of 3 voxel radius, as in the analysis of representational geometry. bsMVPC was performed using a one-nearest neighbor classifier based on correlation distance [12,16]. Each 15 s (6TR) sequence of brain data for an individual was compared to other subjects’ mean responses to that sequence and all other 15 s sequences in the same movie half using a sliding time window, resulting in over 1300 alternative time segments (chance classification accuracy < 0.1%). Classification accuracies in each searchlight were averaged across both halves in each subject before mapping the subject means onto searchlight center voxels on the cortical surface for visualization.

We implemented our methods and ran our analyses in PyMVPA [45](http://www.pymvpa.org) unless otherwise specified. All preprocessing and analyses were carried out on a 64-bit Debian 7.0 (wheezy) system with additional software from NeuroDebian [52](http://neuro.debian.net).

## Acknowledgements

We would like to thank Yaroslav O. Halchenko, Michael Hanke, Sam Nastase, and Nikolaas O. Oosterhof for discussion and software support. This work was supported by grants from the National Institute of Mental Health (R01MH075706) and the National Science Foundation (NSF1129764 and NSF1607845). Resting state and task fMRI data were provided by the Human Connectome Project, WU-Minn Consortium (Principal Investigators: David Van Essen and Kamil Ugurbil; 1U54MH091657) funded by the 16 NIH Institutes and Centers that support the NIH Blueprint for Neuroscience Research; and by the McDonnell Center for Systems Neuroscience at Washington University.

## Supporting information legends

S1 Text. Overview of supplemental data and figures on common model connectome based on responses to a movie.

S2 Fig. Spatial granularity of shared connectivity profiles from the movie data. The common model connectome based on the movie data produced similar results to the common model connectome based on resting state data in terms of ISC of connectivity profiles (Fig 5) and the spatial granularity, as indexed by the PSF of ISCs. (A) Mean PSF slopes in functional ROIs. (B) Mean PSF across all ROIs.

S3 Fig. ISC of representational geometries in the responses to movie time-points. Analysis procedure was identical to Fig 4 with results after RHA included for comparison. (A) ISC of representational geometry in each voxel mapped onto cortical surfaces after anatomical alignment, RHA, and CHA. (B) ISC of representational geometries in 24 functional ROIs and their mean after anatomical alignment and in the common model connectome. Bootstrapped testing showed significantly higher ISCs of representational geometry after both CHA and RHA relative to anatomical alignment in all ROIs, and some differences after CHA and RHA (“-” : CHA<RHA; “ns” : no significant difference between ISCs after CHA and RHA; “+” : CHA>RHA). CHA increased the ISC of representational geometry in all of the ROIs (ROI mean ISC=0.291) relative to anatomical alignment (ISC=0.173) (difference=0.118 [0.103, 0.129]), but the increase is slightly, but significantly, less than that provided by RHA (ISC=0.306) (difference=0.015 [0.009, 0.019]).

S4 Fig. Between-subject classification of movie segments. Analysis procedure was identical to Fig 4 with results after RHA included for comparison. (A) Classification accuracies in each searchlight mapped on cortical surfaces after anatomical alignment, RHA, and CHA. (B) Classification accuracies in 24 ROIs covering visual, auditory, cognitive, and social systems across the cortex and their mean after anatomical alignment, RHA, and CHA. Bootstrapped testing showed significant, six to seven-fold higher accuracies after both the hyperalignment methods relative to anatomical alignment (ROI mean bsMVPC is 1.74%, 12.95%, 9.93% after anatomical alignment, RHA, and CHA, respectively and slightly but significantly higher accuracies after RHA relative to CHA (mean difference = 3.02% [2.52%, 3.40%]). (C) Classification accuracies using information from multiple systems across the whole cortex. Dimensionality of the data is reduced using SVD and classification is performed with different set sizes of top singular vectors. Peak accuracy is reached after 200 dimensions for hyperaligned data and at 50 dimensions for anatomically aligned data. Peak accuracy after RHA is 92.98% and after CHA is 89.61% (mean difference = 3.37% [2.32%, 4.99%]).

S5 Fig. ISC of HCP task activation and contrast maps. Connectivity hyperalignment parameters derived from a session of resting state data were applied to the task maps and the correlation of these maps is computed between each subject and the average of others before and after hyperalignment. Hyperalignment improved correlations on average across all tasks and in all but two (Face-Shapes and Body-Average, labeled ns) individual task maps. The average correlation across task maps increased from 0.58 to 0.65 (mean difference = 0.07 [0.06, 0.08]).

S6 Table. Selected cortical loci implicated in visual, auditory, cognitive, and social functions from Neurosynth.

S7 Table. Task maps used from the HCP data.

## References

1. Biswal B, Yetkin FZ, Haughton VM, Hyde JS. Functional connectivity in the motor cortex of resting human brain using echo-planar MRI. Magn Reson Med. 1995;34: 537–541.

2. Smith SM, Vidaurre D, Beckmann CF, Glasser MF, Jenkinson M, Miller KL, et al. Functional connectomics from resting-state fMRI. Trends in Cognitive Sciences. 2013;17: 666–682.

3. Hutchison RM, Womelsdorf T, Allen EA, Bandettini PA, Calhoun VD, Corbetta M, et al. Dynamic functional connectivity: Promise, issues, and interpretations. NeuroImage. 2013;80: 360–378.

4. Sporns O, Chialvo DR, Kaiser M, Hilgetag CC. Organization, development and function of complex brain networks. Trends in Cognitive Sciences. 2004;8: 418–425.

5. Thomas Yeo BT, Krienen FM, Sepulcre J, Sabuncu MR, Lashkari D, Hollinshead M, et al. The organization of the human cerebral cortex estimated by intrinsic functional connectivity. Journal of Neurophysiology. 2011;106: 1125–1165.

6. Wig GS, Laumann TO, Cohen AL, Power JD, Nelson SM, Glasser MF, et al. Parcellating an Individual Subject’s Cortical and Subcortical Brain Structures Using Snowball Sampling of Resting-State Correlations. Cereb Cortex. 2014;24: 2036–2054.

7. Laumann TO, Gordon EM, Adeyemo B, Snyder AZ, Joo SJ, Chen M-Y, et al. Functional System and Areal Organization of a Highly Sampled Individual Human Brain. Neuron. 2015;87: 657–670.

8. Gordon EM, Laumann TO, Adeyemo B, Petersen SE. Individual Variability of the System-Level Organization of the Human Brain. Cereb Cortex. 2017;27: 386–399.

9. Glasser MF, Coalson TS, Robinson EC, Hacker CD, Harwell J, Yacoub E, et al. A multi-modal parcellation of human cerebral cortex. Nature. 2016;536: 171–178.

10. Gordon EM, Laumann TO, Adeyemo B, Huckins JF, Kelley WM, Petersen SE. Generation and Evaluation of a Cortical Area Parcellation from Resting-State Correlations. Cereb Cortex. 2016;26: 288–303.

11. Cohen AL, Fair DA, Dosenbach NUF, Miezin FM, Dierker D, Van Essen DC, et al. Defining functional areas in individual human brains using resting functional connectivity MRI. NeuroImage. 2008;41: 45–57.

12. Haxby JV, Gobbini MI, Furey ML, Ishai A, Schouten JL, Pietrini P. Distributed and Overlapping Representations of Faces and Objects in Ventral Temporal Cortex. Science. 2001;293: 2425–2430.

13. Haxby JV, Guntupalli JS, Connolly AC, Halchenko YO, Conroy BR, Gobbini MI, et al. A Common, High-Dimensional Model of the Representational Space in Human Ventral Temporal Cortex. Neuron. 2011;72: 404–416.

14. Haxby JV, Connolly AC, Guntupalli JS. Decoding Neural Representational Spaces Using Multivariate Pattern Analysis. Annual Review of Neuroscience. 2014;37: 435–456.

15. Grill-Spector K, Weiner KS. The functional architecture of the ventral temporal cortex and its role in categorization. Nat Rev Neurosci. 2014;15: 536–548.

16. Guntupalli JS, Hanke M, Halchenko YO, Connolly AC, Ramadge PJ, Haxby JV. A Model of Representational Spaces in Human Cortex. Cereb Cortex. 2016;26: 2919–2934.

17. Norman KA, Polyn SM, Detre GJ, Haxby JV. Beyond mind-reading: multi-voxel pattern analysis of fMRI data. Trends in Cognitive Sciences. 2006;10: 424–430.

18. Barch DM, Burgess GC, Harms MP, Petersen SE, Schlaggar BL, Corbetta M, et al. Function in the human connectome: Task-fMRI and individual differences in behavior. NeuroImage. 2013;80: 169–189.

19. Smith SM, Beckmann CF, Andersson J, Auerbach EJ, Bijsterbosch J, Douaud G, et al. Resting-state fMRI in the Human Connectome Project. NeuroImage. 2013;80: 144–168.

20. Van Essen DC, Smith SM, Barch DM, Behrens TEJ, Yacoub E, Ugurbil K. The WU-Minn Human Connectome Project: An overview. NeuroImage. 2013;80: 62–79.

21. Yarkoni T, Poldrack RA, Nichols TE, Essen DCV, Wager TD. Large-scale automated synthesis of human functional neuroimaging data. Nature Methods. 2011;8: 665–670.

22. Robinson EC, Jbabdi S, Glasser MF, Andersson J, Burgess GC, Harms MP, et al. MSM: A new flexible framework for Multimodal Surface Matching. NeuroImage. 2014;100: 414–426.

23. Sabuncu M, Singer BD, Conroy B, Bryan RE, Ramadge PJ, Haxby JV. Function-based intersubject alignment of human cortical anatomy. Cerebral Cortex. 2010;20:130–140.

24. Conroy BR, Singer BD, Haxby JV, Ramadge PR. MRI-Based inter-subject cortical alignment using functional connectivity. In Y Bengio, D Schuurmans, J Lafferty, CKI Williams, A Culotta (eds), Advances in Neural Information Processing Systems 22. 2009: pp 378–386.

25. Conroy BR, Singer BD, Guntupalli JS, Ramadge PR, Haxby JV. Inter-subject alignment of human cortical anatomy using functional connectivity. Neuroimage. 2013;81:400–411.

26. Guntupalli JS, Wheeler KG, Gobbini MI. Disentangling the representation of identity from head view along the human face processing pathway. Cerebral Cortex. 2017;27:46–53.

27. Connolly AC, Sha L, Guntupalli JS, Oosterhof N, Halchenko YO, Nastase SA, et al. How the Human Brain Represents Perceived Dangerousness or “Predacity” of Animals. J Neurosci. 2016;36: 5373–5384.

28. Visconti di Oleggio Castello M, Halchenko YO, Guntupalli JS, Gors JD, Gobbini MI. The neural representation of familiar and unfamiliar faces in the distributed system for face perception. Scientific Reports. 2017;7:12237.

29. Heinzle J, Kahnt T, Haynes J-D. Topographically specific functional connectivity between visual field maps in the human brain. NeuroImage. 2011;56: 1426–1436.

30. Jbabdi S, Sotiropoulos SN, Behrens TE. The topographic connectome. Current Opinion in Neurobiology. 2013;23: 207–215.

31. Haak KV, Marquand AF, Beckmann CF. Connectopic mapping with resting-state fMRI. Neuroimage. 2017;epub ahead of print.

32. Langs G, Sweet A, Lashkari D, Tie Y, Rigolo L, Golby AJ, Golland P. Decoupling function and anatomy in atlases of functional connectivity patterns: Language mapping in tumor patients. Neuroimage. 2014;103:462–475.

33. Langs G, Wang D, Golland P, Mueller S, Pan R, Sabuncu MR, Sun W, Li K, Liu H. Identifying shared networks in individuals by decoupling functional and anatomical variability. Cerebral Cortex. 2016;26:4004–4014.

34. Beckmann CF, Mackay CE, Filippini N, Smith SM. Group comparison of resting-state FMRI data using multi-subject ICA and dual regression. Organization of Human Brain Mapping Abstracts. 2009.

35. Filippini N, MacIntosh BJ, Hough MG, Goodwin GM, Frisoni GB, Smith SM, et al. Distinct patterns of brain activity in young carriers of the APOE-e4 allele. Proceedings of the National Academy of Sciences, USA. 2009;106:7209–7214.

36. Yacoub E, Harel N, Uğurbil K. High-field fMRI unveils orientation columns in humans. PNAS. 2008;105:10607–10612.

37. Nishimoto S, Vu AT, Naselaris T, Benjamini Y, Yu B, Gallant JL. Reconstructing Visual Experiences from Brain Activity Evoked by Natural Movies. Current Biology. 2011;21: 1641–1646.

38. Kanwisher N. Functional specificity in the human brain: A window into the functional architecture of the mind. PNAS. 2010;107: 11163–11170.

39. Freiwald WA, Tsao DY. Functional Compartmentalization and Viewpoint Generalization Within the Macaque Face-Processing System. Science. 2010;330: 845–851.

40. Nastase SA, Connolly AC, Oosterhof NN, Halchenko YO, Guntupalli JS, Visconti di Oleggio Castello M, Gors J, Gobbini MI, Haxby JV. Attention selectively reshapes the geometry of distributed semantic representation. Cereb Cortex. 2017;27: 4277–4291.

41. DiCarlo JJ, Cox DD. Untangling invariant object recognition. Trends in Cognitive Sciences. 2007;11: 333–341.

42. Cox RW. AFNI: software for analysis and visualization of functional magnetic resonance neuroimages. Comput Biomed Res. 1996;29: 162–173.

43. Saad ZS, Reynolds RC, Argall B, Japee S, Cox RW. SUMA: an interface for surface-based intra- and inter-subject analysis with AFNI. IEEE International Symposium on Biomedical Imaging: Nano to Macro, 2004. 2004. Vol. 2; pp. 1510–1513.

44. osterhof NN, Wiestler T, Downing PE, Diedrichsen J. A comparison of volume-based and surface-based multi-voxel pattern analysis. NeuroImage. 2011;56: 593–600

45. Hanke M, Halchenko YO, Sederberg PB, Hanson SJ, Haxby JV, Pollmann S. PyMVPA: A Python toolbox for multivariate pattern analysis of fMRI data. Neuroinformatics. 2009;7: 37–53.

46. Schönemann PH. A generalized solution of the orthogonal procrustes problem. Psychometrika. 1966;31: 1–10.

47. Gower JC. (1975). Generalized procrustes analysis. Psychometrika. 1975;40: 33–51.

48. Chen P-H, Chen J, Yeshurun Y, Hasson U, Haxby JV, Ramadge PJ. A Reduced-dimension fMRI Shared Response Model. Proceedings of the 28th International Conference on Neural Information Processing Systems. Cambridge, MA, USA: MIT Press; 2015. pp. 460–468.

49. Xu H, Lorbert A, Ramadge PJ, Guntupalli JS, Haxby JV. Regularized hyperalignment of multi-set fMRI data. 2012 IEEE Statistical Signal Processing Workshop (SSP). IEEE; 2012. pp. 229–232.

50. Kriegeskorte N, Simmons WK, Bellgowan PSF, Baker CI. Circular analysis in systems neuroscience: the dangers of double dipping. Nat Neurosci. 2009;12: 535–540.

51. Kirby KN, Gerlanc D. BootES: An R package for bootstrap confidence intervals on effect sizes. Behav Res. 2013;45: 905–927.

52. Halchenko YO, Hanke M. Open is Not Enough. Let’s Take the Next Step: An Integrated, Community-Driven Computing Platform for Neuroscience. Front Neuroinform. 2012;6.

